# ESPClust: Unsupervised identification of modifiers for the effect size profile in omics association studies

**DOI:** 10.1101/2024.08.11.607486

**Authors:** Francisco J. Pérez-Reche, Nathan J. Cheetham, Ruth C.E. Bowyer, Ellen J. Thompson, Francesca Tettamanzi, Cristina Menni, Claire J. Steves

## Abstract

High-throughput omics technologies have revolutionised the identification of associations between individual traits and underlying biological characteristics, but still use ‘one effect-size fits all’ approaches. While covariates are often used, their potential as effect modifiers often remains unexplored. To bridge this gap, we introduce ESPClust, a novel unsupervised method designed to identify covariates that modify the effect size of associations between sets of omics variables and outcomes. By extending the concept of moderators to encompass multiple exposures, ESPClust analyses the effect size profile (ESP) to identify regions in covariate space with different ESP, enabling the discovery of subpopulations with distinct associations. Applying ESPClust to insulin resistance and COVID-19 symptom manifestation, we demonstrate its versatility and ability to uncover nuanced effect size modifications that traditional analyses may overlook. By integrating information from multiple exposures, ESPClust identifies effect size modifiers in datasets that are too small for traditional univariate stratified analyses. This method provides a robust framework for understanding complex omics data and holds promise for personalised medicine.

## Introduction

Rapidly expanding high-throughput technologies offer an unprecedented ability to identify associations between observed traits of individuals and biological endpoints via characterisation of various ‘omics’ data. ‘Omics’ represents a range of disciplines including genomics, proteomics, metagenomics and allows elucidation of mechanisms and processed underpinning health and disease states ^1–5^.

Associations between omics variables and a given phenotypic outcome are often influenced by various covariates, including demographics (such as age, sex, body mass index -BMI-, ethnicity), biological diversity within samples^6^ or technical variation in the timing of collection, collection method and processing of samples^7^. It is standard practice to account for the potential confounding effects of such covariates when analysing associations between omics variables and outcomes.

Covariates can also act as moderators which alter the effect size of associations^8^. This possibility, however, is not systematically considered in omics association studies. Neglecting moderators not only risks skewing effect size estimates but also disregards vital heterogeneity linked to covariates, which reflects diversity within human populations. Recognising such diversity can prove invaluable in identifying target subpopulations for maximising intervention effectiveness or delineating thresholds that differentiate groups based on distinct characteristics. Understanding how effects are different within different subpopulations is the cornerstone of developing personalised medicine.

Here, we present ESPClust, an unsupervised method to identify covariates that modify the effect size of association between a set of omics variables and an outcome, which can be used in relatively small sample sizes for discovery science. This method extends the concept of moderators, which typically apply to the relationship between a single exposure and an outcome, to encompass multiple exposures simultaneously. It does so by analysing the effect size profile (ESP), a collection of effect sizes representing the connections between various omics variables and the outcome. The method divides the space of covariates into regions of approximately homogeneous ESP, defining clusters of individuals who share similar associations between their omics profile and the outcome. In essence, ESPClust extends the concept of effect modification to consider modification of the ESP, i.e. modification of the joint effect size of multiple exposures.

ESPClust is versatile and can be readily employed to explore connections between different omics datasets and outcomes. The research questions we address here are chosen to illustrate the ability of the method to a) make new discoveries in a highly researched area and b) identify completely novel findings in an emerging disease. For the former, we investigated the correlation between blood metabolomics and insulin resistance, an area which has already been heavily researched^9–11^, providing a valuable context for our new findings. For the latter, we explored whether and how pre-pandemic blood metabolomics, reflecting pre-infection metabolism, influenced whether someone would become symptomatic after SARS-CoV-2 infection. Although numerous studies have analysed potential relationships between COVID-19 and blood metabolomics, they often concentrate on blood samples collected post-infection or use infection severity (such as hospitalisation) as the primary outcome^12–16^. By considering a new question, we showcase the ability of ESPClust to analyse previously unexplored associations.

## Results

### Overview of the ESPClust method

Given a set of *M* omics variables {*X*_1_, *X*_2_, …, *X*_*M*_}, we define the effect size profile (ESP) as the set of effect sizes {*e*_1_, *e*_2_, …, *e*_*M*_} giving the individual association of each omics variable and the outcome (Fig. 1a). Suppose that each effect size *e*_*m*_ in the ESP can depend on *J* covariates, {*Z*_1_, *Z*_2_, …, *Z*_*J*_}, which act as effect modifiers. ESPClust generalizes the idea of univariate effect modification^8,17^ to modification of the whole ESP (Fig. 1b). The main aim of the method is to identify regions in the covariate space where the ESP can be considered homogeneous. Analyses that ignore effect size modification should be appropriate within each of these regions. In contrast, assuming homogeneity for the ESP across different regions would not be justified.

**Figure 1.**
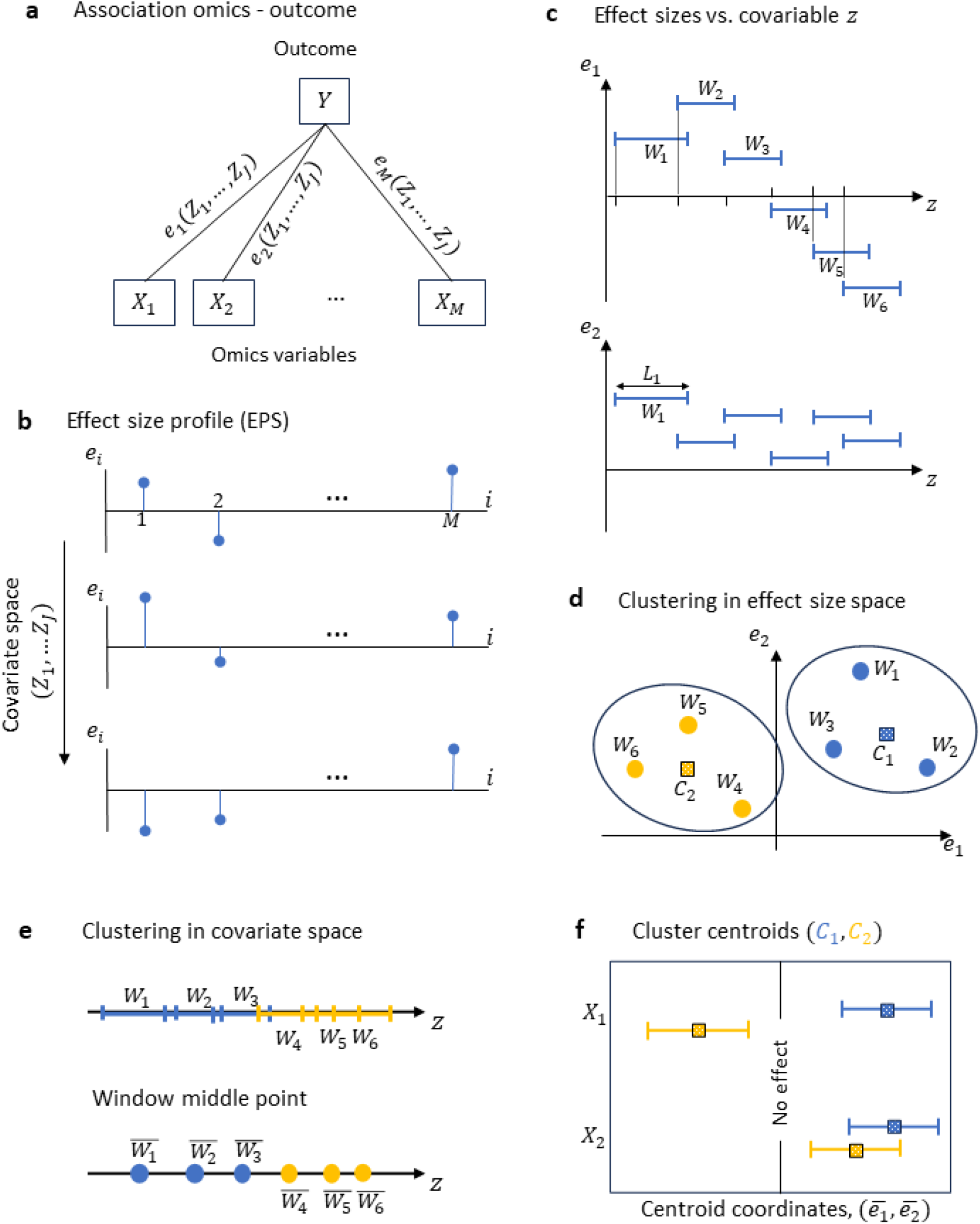
ESPClust method to identify regions in the covariate space with similar effect size profile. (a) Association between *M*omics variables (exposures) and an outcome *Y* in terms of pair-wise effect sizes {*e*_1_, *e*_2_, …, *e*_*M*_} that may depend on *J* covariates. (b) Schematic representation of the effect size profile dependence on the covariates. Panels (c)-(f) illustrate the method for a simple case with two omics variables, {*X*_1_, *X*_2_}, which depend on a single continuous covariate, *z*. (c) The effect size for each omics variable is calculated within 6 windows, 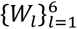, of lengths 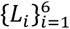 that cover the values taken by the covariate *z*. (d) Clustering of the windows in the effect size space. Windows within a cluster have a similar effect size profile. *C*_1_ and *C*_2_ are the cluster centroids. (e) Window clusters in the covariate space defining regions shown with segments (top) or window midpoints (bottom). (f) Coordinates of the cluster centroids summarising the effect of the covariate *z* on the association profile of each omics variable with the outcome.

ESPClust consists of three basic steps:

1. Evaluation of the ESP as a function of the potential effect modifiers, {*Z*_1_, *Z*_2_, …, *Z*_*J*_}. For a discrete covariate *Z* such as sex or smoking status, simply requires estimating the ESP separately for each value of *Z*. To deal with continuous covariates, we evaluate the ESP within a set of windows 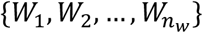 which cover the region of the covariate space spanned by the data. For a single covariate (i.e. for *J* = 1), windows are one-dimensional segments of lengths 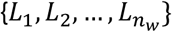 (Fig. 1c). This procedure ultimately approximates the dependence of ESP on the covariates {*Z*_1_, *Z*_2_, …, *Z*_*J*_} by window-dependent effect size spectra (e.g. the ESP corresponding to window *W*_1_ is {*e*_1_(*W*_1_), *e*_2_(*W*_1_), …, *e*_*M*_(*W*_1_)}).
2. Clustering of windows with similar ESP. The ESP for a given window is used as a vector of features that allows windows to be clustered in groups with similar ESP. Fig. 1d illustrates the concept: Windows *W*_1_, *W*_2_ and *W*_3_ used in Fig. 1c form a cluster where both *e*_1_ and *e*_2_ show a positive association across all three windows. In contrast, windows *W*_4_, *W*_5_ and *W*_6_ form a separate cluster where the values of *e*_1_ are negative and those of *e*_2_ are positive. The number of clusters identified by ESPClust can be manually set to any positive integer. However, we will present results in which the number of clusters represents the most frequently occurring value among four clustering indices: Calinski-Harabasz^18^, Davies-Bouldin^19^, silhouette^20^ and elbow^21^ (see Methods). If there is no repeated number across the four indices or in case of a tie, we will use the smallest number of clusters. Agglomerative clustering will be used throughout the article.
3. Identification of regions in the covariate space with homogeneous ESP using the window clusters obtained in step 2. Figure 1e illustrates the process for a case with a single effect modifier. In this example, the covariate space is split into two regions with “small” and “large” *z*. Since the windows covering the covariate space can overlap, hard clustering of windows in the effect size space results in fuzzy clustering in the covariate space^22^. For instance, in Fig. 1e, segments overlap, allowing a specific value of the modifier to belong to multiple windows simultaneously. This approach acknowledges that individuals sharing a common modifier (e.g., age) may exhibit varied associations between omics variables and outcomes. Nonetheless, we employ the midpoint of each window, 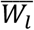, to provide a clearer visualization of regions characterized by homogeneous ESP.

The centroids of the clusters serve as summaries of the ESP within each region, aiding in the interpretation of how covariates influence the relationship between each omics variable and the outcome. In Fig. 1f, the coordinates of the centroids and their dispersion are displayed. In this instance, the influence of the covariate *z* on the association between the pair (*X*_1_, *Y*) is significantly greater than that on the pair (*X*_2_, *Y*). This is evident from the larger difference between the centroids *C*_1_ and *C*_2_ for *X*_1_ compared to those for *X*_2_. Ultimately, this discrepancy reflects the stronger dependency of *e*_1_ on *z*, as illustrated in Fig. 1c.

### Association between insulin resistance and serum metabolomics

We employed ESPClust to investigate the impact of BMI, sex, and gut microbiome gene richness (i.e., the number of unique microbial genes) on the relationship between serum metabolites and insulin resistance (HOMA-IR) among 275 non-diabetic individuals from the Danish MetaHIT study ^11,23,24^ (see Supplementary Table S1). Results will be presented for two examples using different metabolomic variables as exposures.

The first example utilised 94 polar metabolites^11^. To assess the effect sizes of step 1 in ESPClust for each sex, univariate linear regression was employed within predefined windows with several dimensions. Fig. 2 shows the results corresponding to windows determined by sliding a rectangular frame with dimensions (*L*_*BMI*_, *L*_*g*.*rich*._) = (8 kg/m^2^, 0.2*e*6) at steps (Δ_*BMI*_, Δ_*g*.*rich*._) = (1 kg/m^2^, 0.05*e*6), as illustrated by the grey rectangle in Fig. 2c. Only windows containing more than 10 observations (*n* > 10) were considered. Within each window, the effect size for every metabolite was calculated as the slope of a linear regression model, adjusting for both BMI and gene richness. Fig. 2a shows the dependence of the effect size on each of the covariates for two metabolites which exemplify different levels of heterogeneity: the association between aminomalonic acid and insulin resistance exhibits more pronounced variation with the covariates compared to that of decanoic acid.

**Figure 2.**
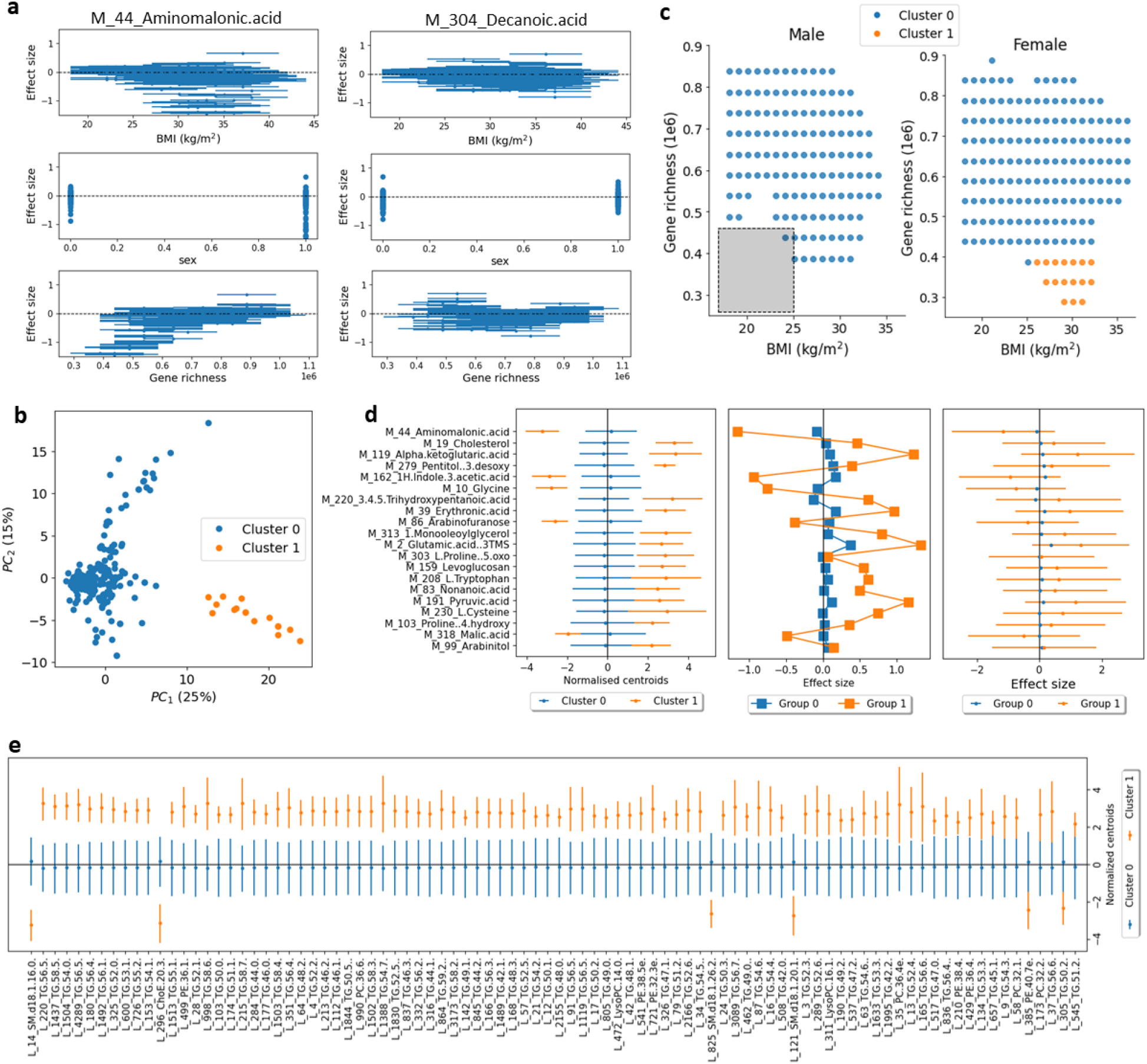
Application of ESPClust to study the association between insulin resistance and (a-d) 94 serum metabolites or (e) 289 molecular lipids. (a) Example of the effect size within windows for aminomalonic acid and decanoic acid in the covariate space (BMI, sex, gene richness). The error bars in the plots for BMI and gene richness indicate the size of the window used to cover these covariates. (b) Visualisation of two clusters with different ESP using the first two principal components of the windows effect sizes. (c) Clusters in the covariate space separately shown for male and female. The symbols indicate the middle point of the windows used to estimate the effect sizes. The size of the window used to calculate effect sizes for fixed sex is shown by a grey rectangle. (d) The left panel shows the coordinates of the cluster centroids corresponding to the 20 metabolites that differ the most between clusters. The error bars indicate 1.96*SD*, where *SD* is the standard deviation of the centroid coordinates. The central panel shows the effect size for the same metabolites for two groups of individuals representing the two identified clusters. The third panel shows the same effect sizes as in the second panel but the 95% confidence intervals for the effect sizes are shown with error bars. (e) Centroid coordinates for the 90 lipids which differ the most between the two clusters.

A split of the windows into two clusters is the most frequent optimal across different clustering indices (Supplementary Fig. S1). This also looks appropriate when projecting the window effect sizes onto the two first principal components (Fig. 2b). Fig. 2c shows that cluster 1 comprises female individuals with high BMI and low gene richness. It cannot be established whether cluster 1 extends to the region of high BMI and low gene richness for males due to insufficient data in this region for this group. Similarly, there is insufficient data to ascertain whether cluster 1 extends to males within the low gene richness range.

In Fig. 2d (left), the centroid coordinates corresponding to the 20 metabolites with the most significant differences between clusters are shown. This highlights that, in addition to differing in ESP collectively, distinct clusters also diverge at the level of specific individual metabolites (such as cholesterol or glycine).

Previous studies have suggested that elevated levels of cholesterol are linked with obesity (i.e. high BMI), increased insulin resistance^25^, and reduced gene richness^26^. Our findings suggest that BMI and insulin resistance may not only confound the association between cholesterol and insulin resistance, but also act as effect modifiers, amplifying the positive association in cluster 1 (i.e. among individuals with high BMI and low gene diversity). Similarly, ESPClust indicates that the negative association between glycine levels and insulin resistance^27^ is moderated by BMI and gene diversity.

To test the sensitivity of ESPClust to the specific dimensions of the windows used, we replicated the earlier analysis using gliding windows of varying sizes. Employing gliding rectangles with dimensions (*L*_*BMI*_, *L*_*g*.*rich*._) = (5 kg/m^2^, 0.2*e*6) once again yields two clusters, segregating individuals with high BMI and low gene richness from the reminder (Supplementary Fig. S2). The sampling of the covariate space becomes more restricted, as indicated by the emergence of gaps associated with windows containing insufficient data (n≤10) to compute effect sizes. For windows of dimensions (*L*_*BMI*_, *L*_*g*.*rich*._) = (8 kg/m^2^, 0.1*e*6), only the region of high gene richness can be explored (Supplementary Fig. S3), since none of the windows in the low gene richness region contain enough observations.

To compare the results of ESPClust with traditional univariate methods using stratification, we run two analyses. In the first analysis, we categorised individuals into two groups representing the clusters identified by ESPClust. Cluster 1 was represented by individuals with BMI ≥ 26 kg/m^2^ and gene richness^26,28^ ≤ 480,000; cluster 0 was represented by the rest of individuals.

Fig. 2d (center) shows that the effect size for the most distinct metabolites exhibits trends similar to those of the centroid coordinates (Fig. 2d (left)). However, the differences in effect sizes do not achieve statistical significance, as evidenced by the overlap of error bars in Fig. 2d (right). This demonstrates that ESPClust can identify clusters with varying ESP levels that may not be discernible combining traditional stratification and univariate analysis. Indeed, ESPClust may detect collective differences between effect size profiles that may not be prominent at the level of individual omics variables unless large datasets are used to enhance the power of univariate analysis.

For the second stratification analysis, we used ESPClust with strata of dimensions (*L*_*BMI*_, *L*_*g*.*rich*._) = (8 kg/m^2^, 0.2*e*6) built with non-overlapping windows gliding at steps that match their size, i.e. (Δ_*BMI*_, Δ_*g*.*rich*._) = (*L*_*BMI*_, *L*_*g*.*rich*._). The results are significantly less informative than those shown in Fig. 2 for overlapping windows. ESPClust gives seven clusters of strata as the most frequent optimal (Supplementary Fig. 4S). Given the small number of strata, this suggests overfitting. When forcing a split of strata into two clusters, ESPClust identifies a cluster for females with low gene richness and high BMI (Supplementary Fig. 5S), analogous to cluster 1 in Fig. 2. However, this cluster consists of a single stratum and does not provide a precise separation between groups in the covariate space.

As a second example within the context of insulin resistance, we utilised ESPClust with 289 lipids from the Danish MetaHIT study as exposures^11^. Our findings mirrored those obtained with 94 metabolites. Employing gliding windows of size (*L*_*BMI*_, *L*_*g*.*rich*._) = (8 kg/m^2^, 0.2*e*6) resulted in a division of the covariate space into two clusters, identical to those depicted in Fig. 2c for the 94 metabolites dataset. Similarly, results for windows of other sizes were consistent with those described above for the 94 metabolites.

The coordinates of the cluster centroids for the lipidomic dataset indicate a higher effect size within cluster 1 for numerous triglycerides (e.g. TG(56:5)) and some glycerophosphoethanolamines (e.g. PE(36:1)) (Fig. 2e). Previous studies have established that elevated levels of triglycerides are linked with obesity, low gene richness, and metabolic disorders such as insulin resistance^28,29^. Our findings suggest that the positive association between triglycerides and insulin resistance is particularly strengthened in regions characterized by high BMI and low gene richness.

In Fig. 2e, three sphingomyelins exhibit a diminished effect size within cluster 1, with SM(d18:1/16:0) (C16 Sphingomyelin) being the most prominent effect modifier. The interpretation of such a negative association is not clear, as previous research has reported a positive correlation between endogenous sphingomyelins and insulin resistance^30^. Conversely, exogenous dietary sphingomyelins were found to be negatively associated with insulin resistance and obesity^31^.

### Association between the COVID-19 symptoms manifestation and pre-pandemic serum metabolites

We utilized ESPClust to explore the potential of BMI, sex, and age as modifiers for the association between metabolomic variables collected before COVID-19 infection and the manifestation of COVID-19 symptoms (i.e. symptomatic or asymptomatic). The symptom status was derived from self-reported symptoms^32^ in TwinsUK COVID-19 questionnaires^33^, administered between July 2020 and February 2022, and serology data^34^ (see Methods).

We will illustrate the performance of ESPClust for exposures taken from two different metabolomics datasets (see Supplementary Table S2).

The first example is based on 221 biomarkers obtained from serum of 680 participants of the TwinsUK cohort study^35^ using Nuclear Magnetic Resonance ^36^ (NMR). The ESP was estimated through univariate logistic regression within a series of windows defined by sliding rectangles of various sizes. Only windows containing more than 25 observations (n>25) were considered, ensuring a robust representation of both symptomatic and asymptomatic phenotypes within each window.

Defining windows with gliding rectangles of size (*L*_*BMI*_, *L*_*age*_) = (8 kg/m^2^, 21 yr) results in three clusters (Fig. 3a). Cluster 2 exclusively comprises male individuals (Fig. 3b-left); clusters 0 and 1 are predominantly found within the female covariate subspace (Fig. 3b-right). We infer that sex acts as a modifier for the EPS characterising the association between NMR biomarkers and the onset of COVID-19 symptoms. However, due to the limited data coverage for male individuals, this conclusion is only applicable to males with relatively low BMI (<29 kg/m2) and aged over 35 years.

**Figure 3.**
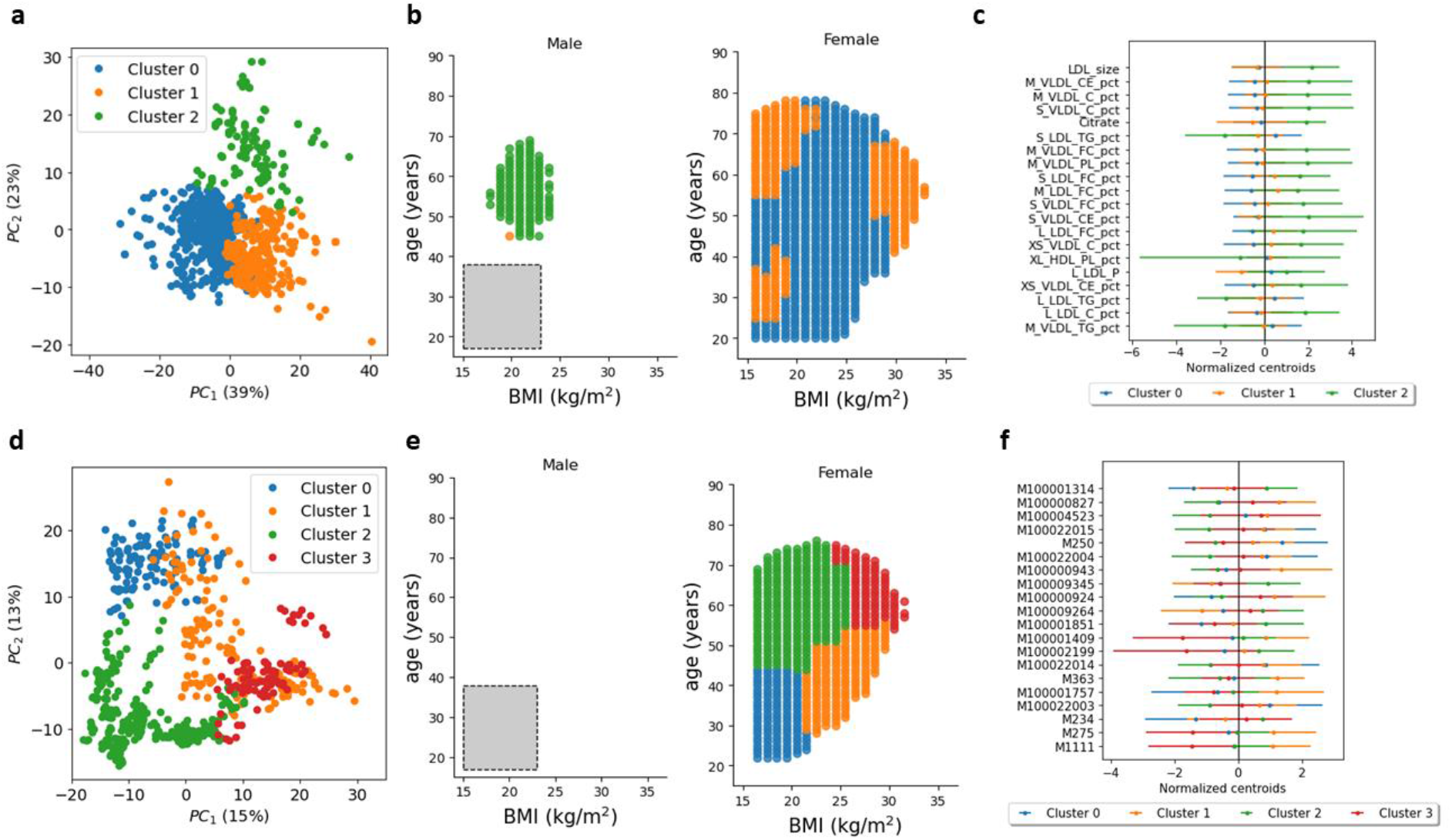
Results obtained by employing EPSClust to investigate the potential role of BMI, sex and age on the association between COVID-19 symptoms manifestation and serum metabolomics. Panels (a-c) show findings from ESPClust analysis using 221 NMR blood biomarkers as exposures, while panels (d-f) present analogous results obtained using 774 LC-MS blood metabolites. (a,d) Visualisation of the clusters for the ESP, using the first two principal components of the window effect sizes. (b,e) Clusters in the covariate space separately shown for male and female individuals. The symbols indicate the middle point of the windows used to estimate effect sizes. The size of the gliding window used to calculate effect sizes for fixed sex is represented by a grey rectangle in the panel for male individuals. (c,f) Coordinates of the cluster centroids corresponding to the 20 metabolites that differ the most between clusters. The error bars indicate 1.96*SD*, where *SD* is the standard deviation of the centroid coordinates.

The differences between clusters 0 and 1, as depicted in Fig. 2b-right, are not easily interpretable in terms of BMI and age. We note, however, a significant overlap between these clusters in the PCA plot of Fig. 3a, suggesting that the disparities in ESP between these two clusters are minor, and combining them to form a single cluster for female individuals would offer a more parsimonious description.

The coordinates of the centroids for different clusters overlap and no individual metabolite exhibits a clearly distinct effect size across clusters (Fig. 3c). In this application, the differences between the clusters are therefore linked to the EPS as a whole. Nevertheless, there is a notable trend for many LDL and VLDL ratios to show higher effect sizes in males compared to females. A larger dataset might lead to a statistically clearer trend in this direction.

When ESPClust was run with smaller gliding windows of size (*L*_*BMI*_, *L*_*age*_) = (5 kg/m^2^, 21 yr) or (8 kg/m^2^, 11 yr), the explored region in the male subspace shrunk, and the ESP differences between males and females could not be identified (Supplementary Figs. S6 and S7).

The second metabolomics dataset utilised comprises 774 pre-pandemic serum metabolites obtained through liquid chromatography-mass spectrometry ^37,38^ (LC-MS) from 368 TwinsUK participants. Employing gliding windows of size (*L*_*BMI*_, *L*_*age*_) = (8 kg/m^2^, 21 yr) results in four clusters in the female subspace (Fig. 3d); none of the windows contained sufficient data to estimate the ESP in the male subspace (Fig. 3e).

In terms of BMI and age, the split of windows in the female subspace obtained using the 774 LC-MS metabolites (Fig. 3b-right) is more intuitive than that obtained using 221 NMR metabolites (Fig. 3e-right). One possible explanation is that increasing the number of metabolites enhances the resolution of the dependence of the EPS on the covariates. At the level of individual metabolites, however, there is again significant overlap between clusters (Fig. 3f).

## Discussion

We have introduced ESPClust, a flexible method for unsupervised identification of effect size modifiers in omics association studies. The method expands upon the concept of effect size modification, traditionally related to the association between an exposure-outcome pair, to utilise the information provided by a set of effect sizes for the association of multiple omics variables and an outcome. This collection of effect sizes defines the effect size profile, referred to as ESP. ESPClust finds regions in the covariate space with differing ESP, effectively generalising the effect size modification concept for individual omics variables to the concept of ESP modification.

An ESP modifier may function as an effect modifier for individual omics variables, as demonstrated in our analysis of serum metabolites and insulin resistance. However, ESP modification captures phenomena not discernible at the individual omics level, as shown in our COVID-19 symptom phenotype analysis.

ESPClust approximates the dependence of ESP on covariates using a cover of the covariate space consisting of overlapping windows. This concept, rooted in topology^39^, expands upon the traditional disjoint partitioning used in stratified analysis. Overlapping windows offer advantages: they eliminate the arbitrariness in defining strata that may intersect regions with heterogeneous ESPs, such as conventional age groups. They also provide a more detailed description of the dependence of effect sizes on covariates. However, overlapping windows create fuzzy boundaries separating regions with different ESPs.

The dimensions of windows defining a cover are adjustable parameters of ESPClust. Ideally, windows should offer a detailed description of the dependence of ESP on covariates across a wide region, ensuring statistically robust estimates within each window. We suggest running ESPClust with various window settings, as shown in our examples. Future research will explore automatic optimisation of cover configurations. The number of clusters to identify groups with similar ESP is also adjustable. In this study, we used a specific rule for selecting the number of clusters, but exploring different numbers of clusters can be beneficial for identifying suitable divisions.

ESPClust offers considerable potential for advancing personalised medicine by identifying subpopulations with distinct biological responses. By detecting covariate-specific effect size modifications even using relatively small datasets, ESPClust reveals subtle associations that traditional methods may miss. This ability to tailor interventions based on individual biological profiles can enhance treatment efficacy and precision. Consequently, ESPClust facilitates the development of more personalised healthcare strategies, improving patient outcomes and driving progress in the field of personalised medicine.

## Methods

### Clustering methods

In the step 2 of ESPClust, the effect size patterns {*e*_1_(*W*_*l*_), *e*_2_(*W*_*l*_), …, *e*_*M*_(*W*_*l*_)} were normalised before clustering. More explicitly, the effect sizes for a given omics variable, 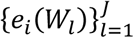, were transformed by subtracting the mean over different windows and dividing by the standard deviation.

To obtain the optimal number of clusters, Calinski-Harabasz, Davies-Bouldin and silhouette clustering measures, the optimal number of clusters clustering corresponds to the number of clusters at the maximum.

The optimal number of clusters for the elbow method was calculated by considering the value of *k* for which the change in slope for the clustering quadratic error ^21^ (also called inertia) is maximal. This effectively identifies the most prominent elbow in the discrete curve obtained by plotting the inertia vs. *k*.

To give the operational definition used by ESPClust, let us denote the inertia of *k* clusters as *I*_*k*_. The slope of the inertia is then *s*_*k*_ = *I*_*k*_ − *I*_*k*−1_ for *k* = 2,3, … From this, one can calculate the relative change of the slope at *k* as follows:

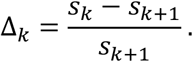

Since the slope *s*_*k*_ is non-positive and increases with *k*, the relative change takes a non-negative value for any *k*. The optimal number of clusters according to the elbow method implemented in ESPClust corresponds to the value of *k* for which Δ_*k*_ is maximum.

### Regression to determine effect sizes within windows

The current implementation of ESPClust uses a linear model to describe the outcome *Y* as a function of an omics variable *X*_*m*_ adjusting for confounding of {*Z*_1_, *Z*_2_, …, *Z*_*J*_}:

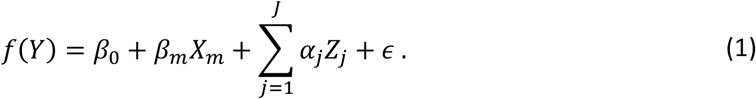

Here, *ϵ* is a normally distributed error. The function *f*(*Y*) depends on the nature of the outcome. For a continuous outcome such as insulin resistence, we used simple linear regression with *f*(*Y*) = *Y*. In this case, the effect sizes are given by the slope coefficient of the model, i.e. *e*_*m*_ = *β*_*m*_. For a binary variable such as the COVID-19 symptomatic/asymptomatic status, we used logistic regression with *f*(*Y*) = logit(*Y*). In this case, the presented effect sizes *e*_*m*_ are the odds ratios given by 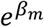.

### Data for the insuline resistance examples

The data used for this example was published by Pedersen et al.^11,24^ who gave a complete description. We restricted our analysis to known metabolites and lipids within these data.

### Data for the COVID-19 symptoms example

#### Study population

The individuals included in this example were selected from the UK Adult Twin Registry (TwinsUK). A study participant was included in the analysis if the following conditions are satisfied: (i) There was pre-pandemic metabolomic data for the participant, (ii) there was information on the presence or absence of COVID-19 symptoms and (iii) there was evidence that the participant was infected by SARS-CoV-2.

#### Exposure variables

The metabolite concentrations of fasting blood samples collected before the COVID-19 pandemic were measured with two different platforms that yielded the two metabolic datasets used in this study. The first dataset was obtained through a high-throughput nuclear magnetic resonance (NMR) platform^36,40^ by Nightingale Health Ltd., Helsinki, Finland. This platform provides the concentration of over 200 circulating metabolic biomarkers including lipids, fatty acids, amino acids, ketone bodies glycolysis related metabolites as well as lipoprotein subclass distribution and particle size. The second dataset (C19-1) was obtained using an untargeted liquid chromatography-mass spectrometry (LC-MS) procedure conducted by Metabolon, Inc., Durham, North Carolina, USA as previously described^37,38^.

#### Outcome variable

Each of the infected cohort participants was classified into one of two classes: asymptomatic or symptomatic. Participants were labelled as asymptomatic if they reported that had not had COVID-19 but there was evidence of SARS-CoV-2 infection. In contrast, participants were assumed to be symptomatic if there was evidence of natural infection and they reported having had COVID-19 and also provided the duration of symptoms (this requirement was imposed to strengthen the evidence that these patients were symptomatic). Information on the symptoms of participants was obtained from three TwinsUK COVID-19 questionnaires^33^ administered in July-August 2020 (Q2), October-November 2020 (Q3) and November 2021-February 2022 (Q4). SARS-CoV-2 infection was assessed using antibody testing data obtained in two rounds that approximately coincide in time with the questionnaires Q2 and Q4. These data were informed by self-reported vaccination status to conclude that there was evidence of SARS-CoV-2 infection for any participant with a positive anti-Nucleocapsid result at any time or a positive anti-Spike result before vaccination^41^.

#### Missing data

Metabolites whose concentration was missing for more than 20% of individuals were discarded. Similarly, individuals who missed more than 20% of the metabolites were also discarded. The remaining missing values for metabolites were imputed using k Nearest Neighbours ^42^ with *k* = 3.

#### Data transformation

Sex was encoded as a numerical variable (0 for male and 1 for female); the rest of variables are intrinsically numerical. Metabolites were individually transformed by adding one and applying the natural logarithm function. All variables were individually standardized by subtracting the mean value and dividing by the standard deviation.

## Data availability

The metabolomic data for the examples on insulin resistance are available at^24^ https://bitbucket.org/hellekp/clinical-micro-meta-integration/src/master/. The data used for the COVID-19 examples are held by the Department of Twin Research at King’s College London. The data can be released to bona fide researchers using our normal procedures overseen by the Wellcome Trust and its guidelines as part of our core funding (https://twinsuk.ac.uk/resources-for-researchers/access-our-data/).

## Code availability

ESPClust software is freely available at https://github.com/fjpreche/ESPClust.git. It can be installed via Python package repositories as `pip install -i https://test.pypi.org/simple/ESPClust==1.0.0`

## Acknowledgements

We acknowledge financial support from the UKRI COVID-19 Longitudinal Health and Wellbeing National Core Study. FJPR acknowledges funding support from a Medical Research Council Fellowship (MR/W021455/1).

## Supplementary Tables

**Table S1.** Demographics relevant to the application of ESPClust to the association between serum metabolomics and insulin resistance. The statistics for insulin resistance, BMI, and gene richness are given as Median (2.5% percentile, 97.5% percentile).

| sex | n | Insulin resistance | BMI (kg/m<sup>2</sup>) | Gene richness |
| --- | --- | --- | --- | --- |
| Male | 125 | 1.8 (0.4,4.9) | 30.9 (20.4,39.9) | 744470 (407391,1013525) |
| Female | 150 | 1.5 (0.5,5.4) | 30.7 (19.6,41.5) | 736092 (378970,958634) |
| All | 275 | 1.7 (0.4,5.2) | 30.7 (20.0,41.2) | 743324 (398365,984865) |

**Table S2.**
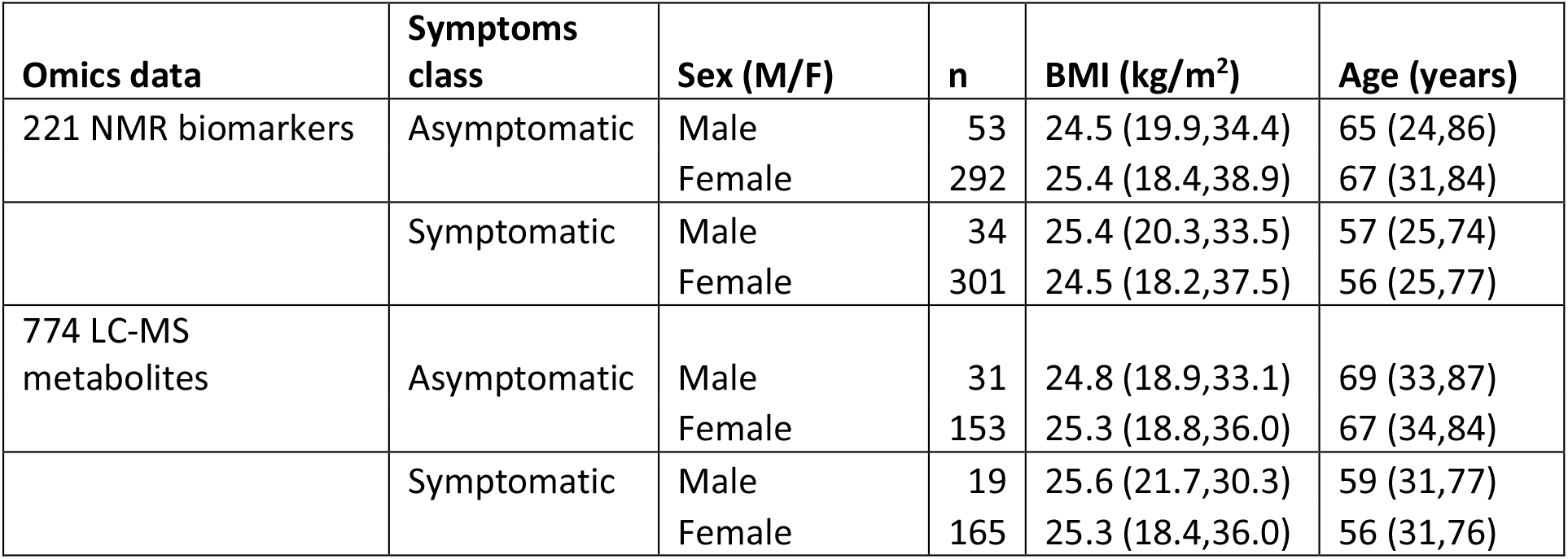
Demographics relevant to the application of ESPClust to the association between serum metabolomics and COVID-19 symptoms manifestation. The statistics for BMI and age are given as Median (2.5% percentile, 97.5% percentile)

| Omics data | Symptoms class | Sex (M/F) | n | BMI (kg/m<sup>2</sup>) | Age (years) |
| --- | --- | --- | --- | --- | --- |
| 221 NMR biomarkers | Asymptomatic | Male | 53 | 24.5 (19.9,34.4) | 65 (24,86) |
|  |  | Female | 292 | 25.4 (18.4,38.9) | 67 (31,84) |
|  | Symptomatic | Male | 34 | 25.4 (20.3,33.5) | 57 (25,74) |
|  |  | Female | 301 | 24.5 (18.2,37.5) | 56 (25,77) |
| 774 LC-MS metabolites | Asymptomatic | Male | 31 | 24.8 (18.9,33.1) | 69 (33,87) |
|  |  | Female | 153 | 25.3 (18.8,36.0) | 67 (34,84) |
|  | Symptomatic | Male | 19 | 25.6 (21.7,30.3) | 59 (31,77) |
|  |  | Female | 165 | 25.3 (18.4,36.0) | 56 (31,76) |

## Supplementary figures

**Figure S4.**
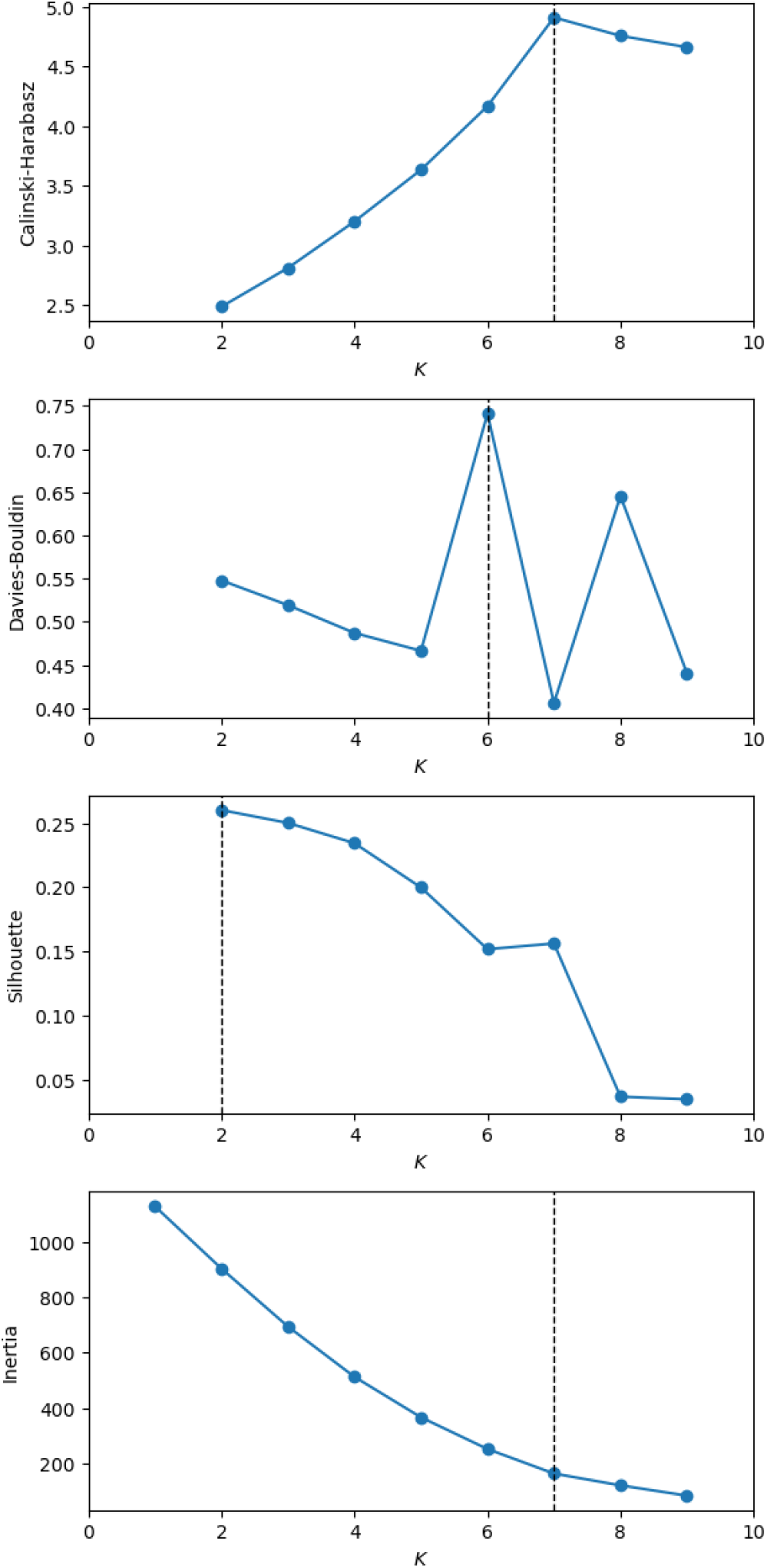
Four clustering measures as a function of the number of clusters for the example on the association of insulin resistance and 94 metabolites using windows of dimensions (*L*_*BMI*_, *L*_*g*.*rich*._) = (8 *kg*/*m*^2^, 0.2*e*6) gliding at steps (*Δ*_*BMI*_, *Δ*_*g*.*rich*._) = (1 *kg*/*m*^2^, 0.05*e*6).

**Figure S5.**
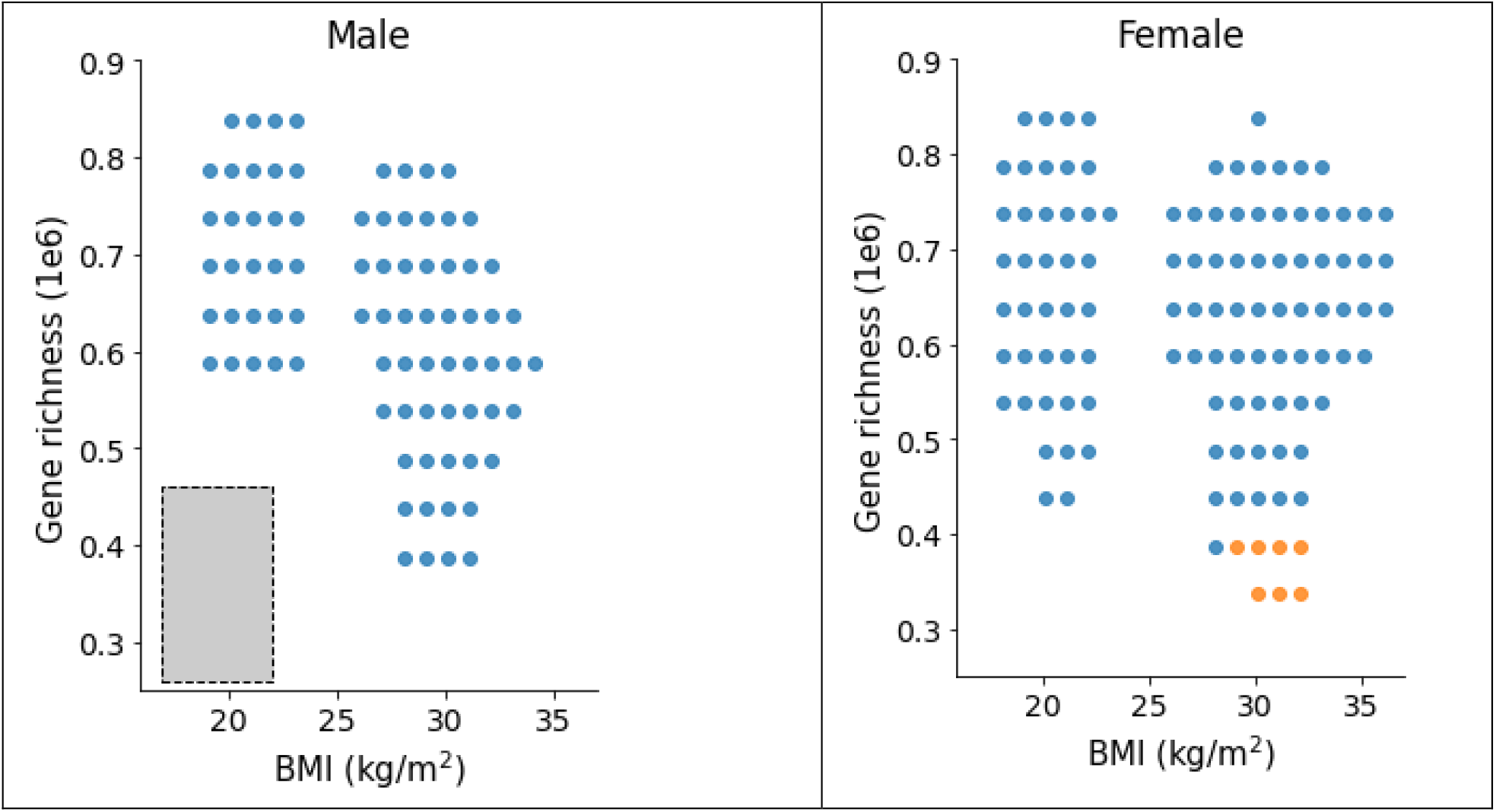
Clusters in the covariate space obtained by applying ESPClust to study the impact of BMI, sex and gene richness on the association of insulin resistance and 94 serum polar metabolites. For given sex, a gliding window of dimensions(*L*_*BMI*_, *L*_*g*.*rich*._) = (5 *kg*/*m*^2^, 0.2*e*6)was used, as marked by the dashed-line rectangle.

**Figure S3.**
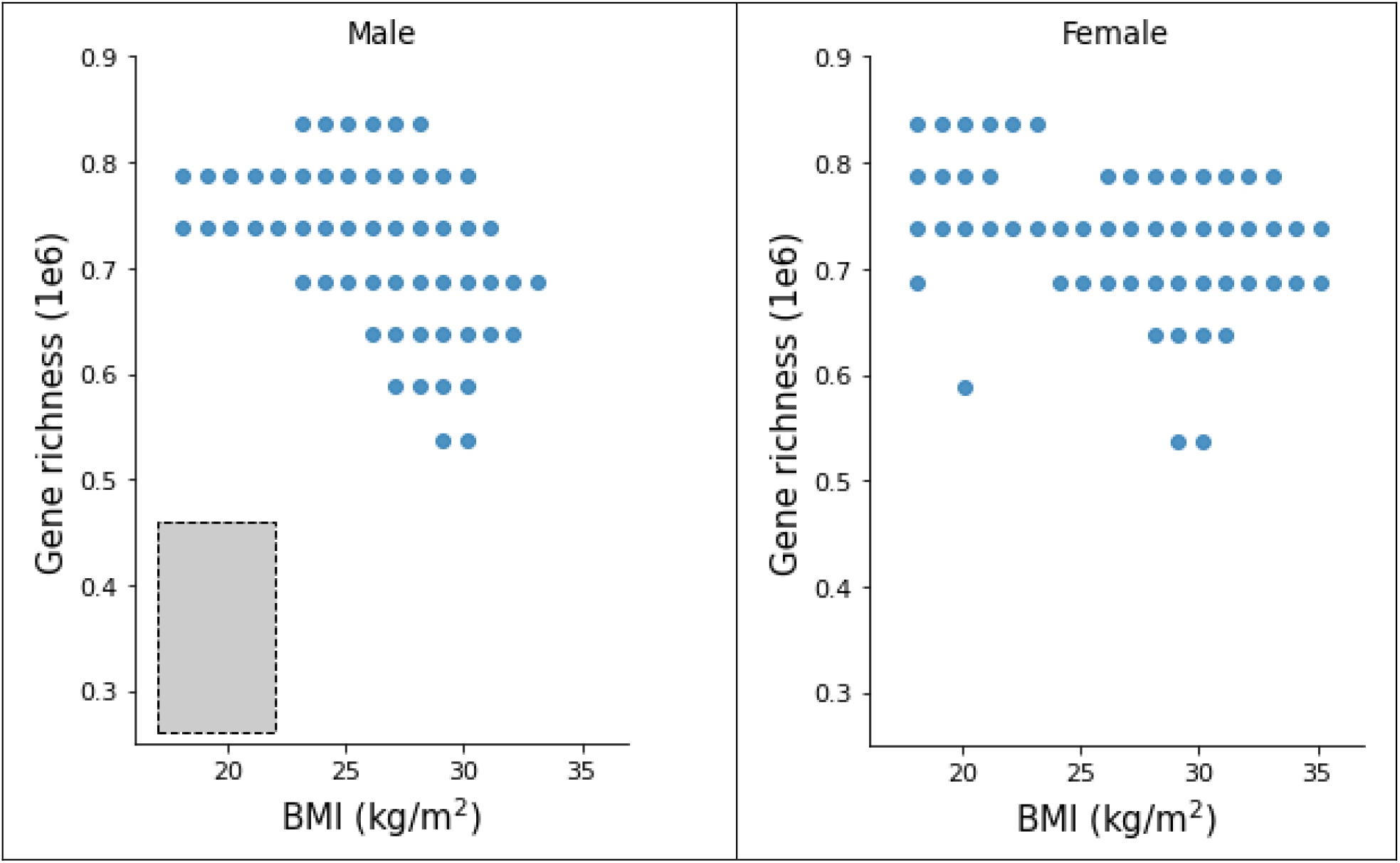
Clusters in the covariate space obtained by applying ESPClust to study the impact of BMI, sex and gene richness on the association of insulin resistance and 94 serum polar metabolites. For given sex, a gliding window of dimensions (*L*_*BMI*_, *L*_*g*.*rich*._) = (8 *kg*/*m*^2^, 0.1*e*6) was used, as marked by the dashed-line rectangle.

**Figure S4.**
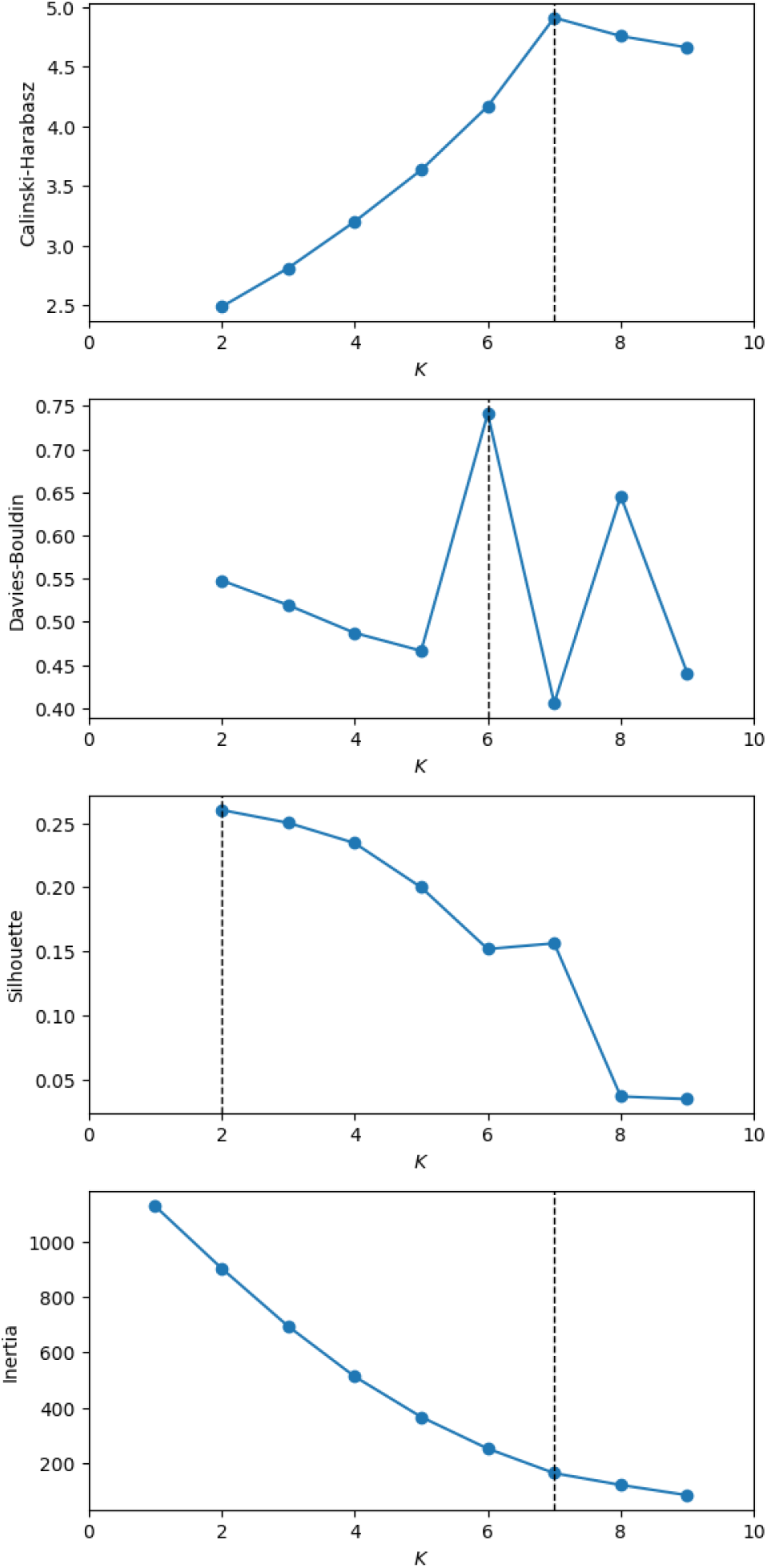
Four clustering measures as a function of the number of clusters for the example on the association of insulin resistance and 94 metabolites using windows of dimensions (*L*_*BMI*_, *L*_*g*.*rich*._) = (8 *kg*/*m*^2^, 0.2*e*6) gliding at steps (*Δ*_*BMI*_, *Δ*_*g*.*rich*._) = (1 *kg*/*m*^2^, 0.05*e*6).

**Figure S5.**
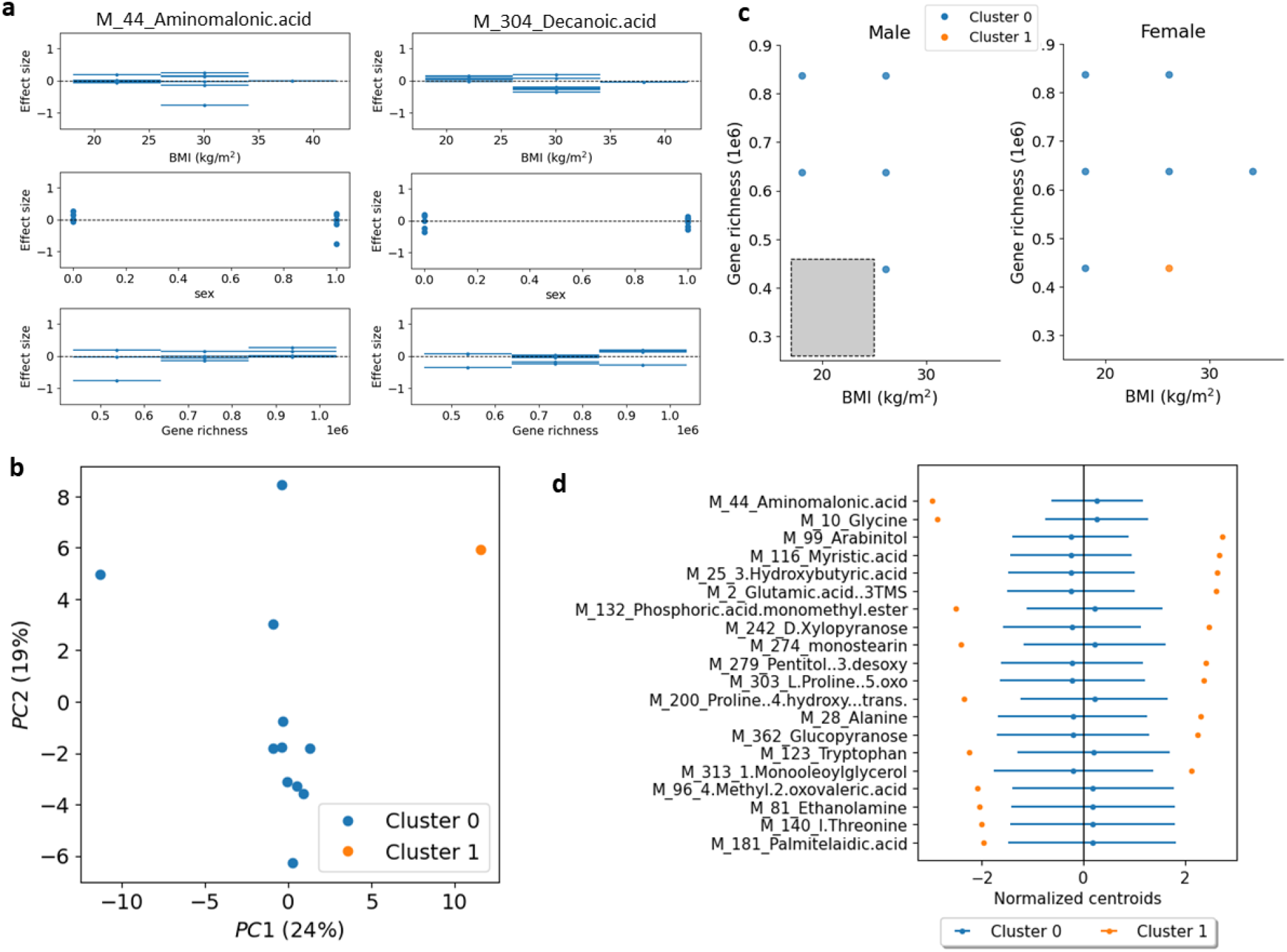
Application of ESPClust to study the association between insulin resistance and 94 serum using *non-overlapping strata* of dimensions (*L*_*BMI*_, *L*_*g*.*rich*._) = (8 *kg*/*m*^2^, 0.2*e*6). (a) Example of the effect size within windows for aminomalonic acid and decanoic acid in the covariate space (BMI, sex, gene richness). The error bars in the plots for BMI and gene richness indicate the size of the window used to cover these covariates. (b) Visualisation of two clusters with different ESP using the first two principal components of the windows effect sizes. (c) Clusters in the covariate space separately shown for male and female. The symbols indicate the middle point of the strata used to estimate the effect sizes. The size of the strata used to calculate effect sizes for fixed sex is shown by a grey rectangle. (d) Coordinates of the cluster centroids corresponding to the 20 metabolites that differ the most between clusters. The error bars indicate 1.96*SD*, where *SD* is the standard deviation of the centroid coordinates.

**Figure S6.**
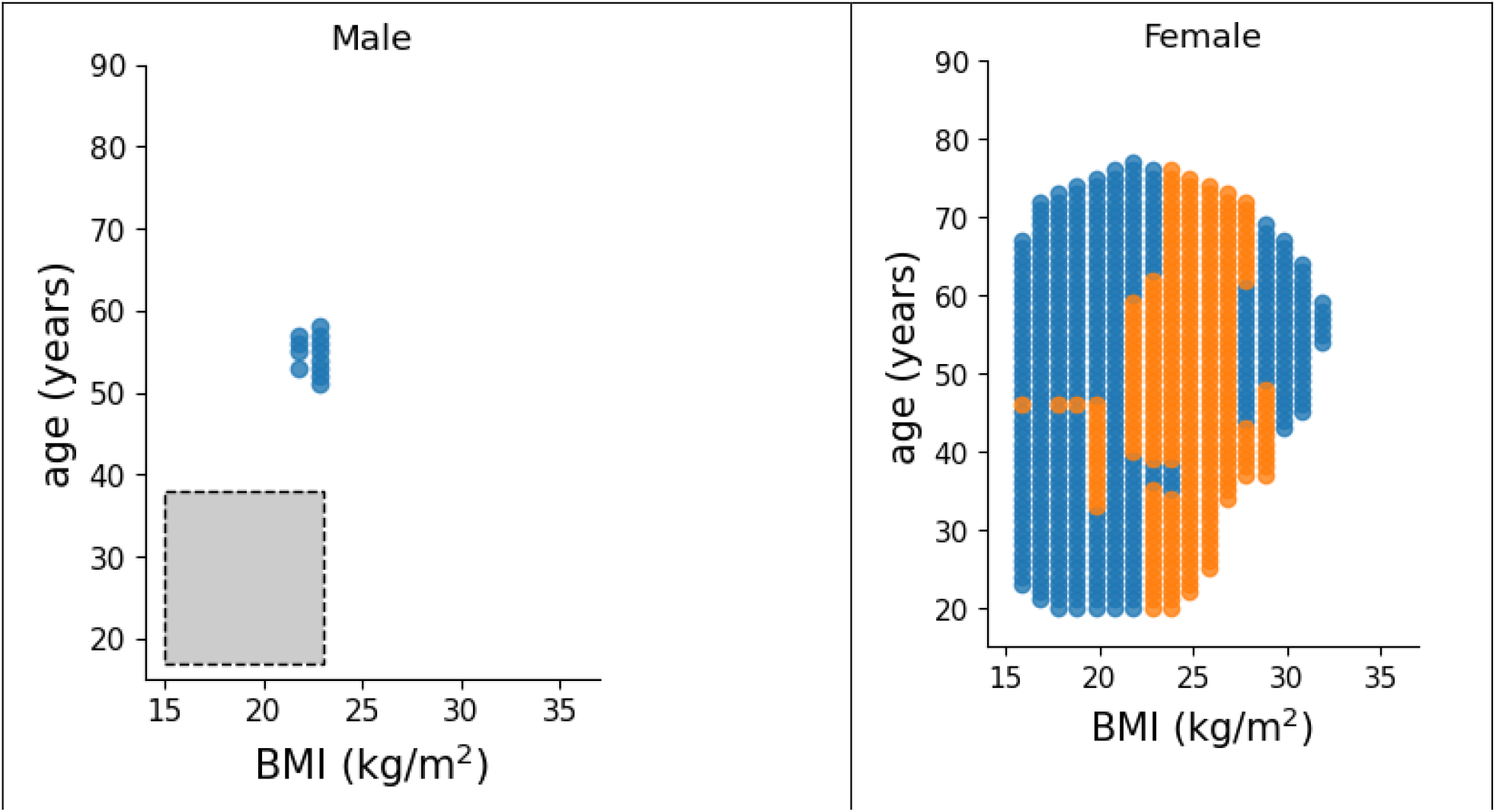
Clusters in the covariate space obtained by applying ESPClust to study the impact of BMI, sex and age on the association of COVID-19 symptoms manifestation and 221 NMR serum metabolites. For given sex, a gliding window of dimensions (*L*_*BMI*_, *L*_*age*_) = (5 *kg*/*m*^2^, 21 *yr*) was used, as marked by the dashed-line rectangle.

**Figure S7.**
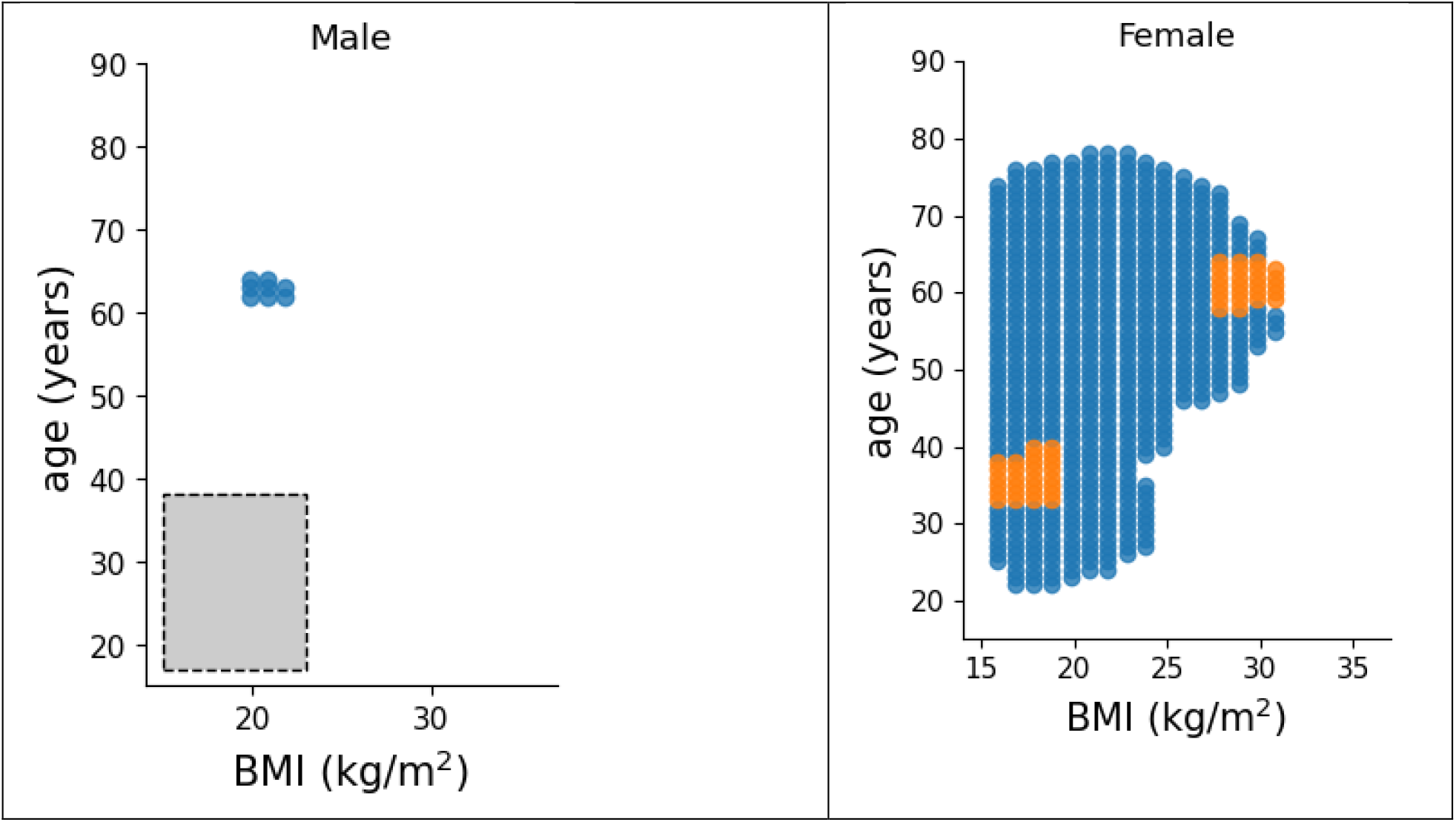
Clusters in the covariate space obtained by applying ESPClust to study the impact of BMI, sex and age on the association of COVID-19 symptoms manifestation and 221 NMR serum metabolites. For given sex, a gliding window of dimensions (*L*_*BMI*_, *L*_*age*_) = (8 *kg*/*m*^2^, 11 *yr*) was used, as marked by the dashed-line rectangle.

## Notes

### Competing Interest Statement

The authors have declared no competing interest.

